# Improved Calreticulin Nanobody by Framework Engineering

**DOI:** 10.64898/2026.06.11.731675

**Authors:** Logan Mavar, Mikhail Pavlenok, Avraneel Paul, Lucinda Hall, Benjamin M. Larimer, Michael Niederweis

## Abstract

Calreticulin is an emerging cancer biomarker, but current detection methods rely on expensive monoclonal antibodies that suffer from inefficient protein production, pharmacokinetic challenges and poor tissue penetration. Cal3, a calreticulin-specific nanobody, was constructed by replacing the complimentary determining region 2 (CDR2) of a soluble, clinically validated nanobody with a calreticulin-specific CDR2 isolated from a phage display library. However, the poor solubility and low yield of Cal3 limit its usefulness. In this study, we engineered CALR-Nb02 by adapting the core of Cal3 to a partial consensus framework sequence of stable nanobodies. CALR-Nb02 was purified with a 240-fold higher yield as a predominantly monomeric, soluble protein that exhibits an increased thermal stability and a higher calreticulin binding affinity (KD: 25–50 nM) compared with Cal3. These results reveal a strategy for quickly altering the specificity of a stable nanobody, and provide an improved calreticulin-binding reagent for future diagnostic, imaging, and therapeutic applications.

## INTRODUCTION

Biofluid-based biomarkers have transformed diagnosis, research and care of many diseases including neurodegenerative dementias (Zetterberg & Bendlin, 2026), multiple sclerosis (Bader *et al*., 2026) and cancer (Passaro *et al*., 2024), triggering intensive research in biomarker discovery (Bader *et al*., 2026, Krusen *et al*., 2026). However, the translation of protein biomarkers into useful clinical tools requires target-specific, high affinity binding reagents (Passaro *et al*., 2024). Currently antibodies are widely used for protein detection in conventional detection assays, but their large size complicates protein production and engineering, impairs tissue penetration and causes other pharmacokinetic challenges (Tang & Cao, 2021, Arbabi-Ghahroudi, 2017, Muyldermans, 2013, Schumacher *et al*., 2018). These limitations have motivated the development of alternative binding reagents that can be produced recombinantly and adapted for diagnostic, imaging, and therapeutic applications.

Nanobodies, single-domain heavy-chain–only antibodies derived from camelids, provide key advantages over mammalian antibodies for biomarker recognition (Muyldermans, 2013). Their small size, single-domain structure, and stability facilitate recombinant production in *E. coli*, enabling scalable and cost-effective manufacturing (Arbabi-Ghahroudi, 2017). Nanobodies can easily be modified for site-specific conjugation to imaging agents or therapeutic payloads (Schumacher *et al*., 2018). Their compact structure can also improve access to specific epitopes, supporting applications in targeted delivery of therapeutic agents. However, despite their reputation for stability and solubility, individual nanobodies can vary substantially in expression yield, folding efficiency, aggregation behavior, and thermal stability (Dingus *et al*., 2022, Kunz *et al*., 2017, Kunz *et al*., 2018). These properties are critical because poorly soluble nanobodies are difficult to produce in large quantities and may contain only a small fraction of functional protein. Therefore, improving the biochemical and biophysical properties of nanobodies is an important step in developing them into useful binding reagents.

Calreticulin is a multifunctional protein that has emerged as a clinically relevant biomarker for multiple cancers (Fucikova *et al*., 2021). Although originally characterized as an endoplasmic reticulum chaperone involved in protein quality control and calcium homeostasis, calreticulin also has roles outside the endoplasmic reticulum. In response to cellular stress, including chemotherapy-induced immunogenic cell death, calreticulin translocates to the cell surface, where it promotes immune recognition and uptake of malignant cells by antigen-presenting cells (Obeid *et al*., 2007). Recent studies have also shown that macrophages secrete a C-terminally cleaved form of calreticulin that decorates target cells and serves as a pro-phagocytic signal (Banuelos *et al*., 2025). Surface-exposed and soluble calreticulin have been associated with cancer prognosis, immune response, and treatment outcome, although the biological implications differ depending on disease context and calreticulin localization (Fucikova *et al*., 2021, Kepp *et al*., 2020). Accordingly, calreticulin has been investigated as a biomarker for multiple cancers, including ovarian cancer, multiple myeloma, triple-negative breast cancer, chemotherapy response, and immunotherapy monitoring (Ak *et al*., 2025, Cao *et al*., 2025, Kielbik *et al*., 2021, Serrano Del Valle *et al*., 2022, Vaes *et al*., 2021, Zhang *et al*., 2021). The findings of these studies establish that calreticulin participates in extracellular and immune-related processes that support its relevance as a cancer biomarker.

Despite the growing interest in calreticulin as a biomarker, tools for its detection remain limited. Cal3 Nb, a previously reported calreticulin-binding nanobody, was engineered by replacing the complimentary determining region 2 (CDR2) of the soluble, clinically validated 2Rs15d nanobody with a calreticulin-specific CDR2 isolated from a synthetic phage display library of randomized CDR2 sequences (D’Huyvetter *et al*., 2017, Hall *et al*., 2025). While Cal3 Nb binds calreticulin with nanomolar affinity, its poor solubility and strong tendency to aggregate limit its usefulness in research and clinical applications.

In this study, we improved Cal3 Nb by adapting its core to a partial consensus sequence identified in highly stable and soluble nanobodies to improve recombinant expression without disrupting antigen recognition (Dingus *et al*., 2022). We engineered two framework-modified nanobodies derived from Cal3 Nb and produced them in the cytoplasm of *E. coli* with 240- and 720-fold improved yields. This approach produced CALR-Nb02, an engineered calreticulin-specific nanobody with increased thermal stability and a higher calreticulin binding affinity compared with the parent nanobody Cal3 Nb. These findings show that partial consensus framework engineering can rescue a poorly soluble but antigen-specific nanobody and not only reveal a strategy for quickly altering the specificity of a stable nanobody but also provide an improved calreticulin-binding reagent for future diagnostic, imaging, and therapeutic applications.

## RESULTS

### Production of the Cal3 Nanobody in *E. coli*

In a previous study, the complementarity-determining region (CDR) 2 of the clinically validated HER2-specific nanobody 2Rs15d was randomized in a synthetic phage display library. Screening for the best calreticulin binders resulted in the high-affinity Calreticulin-specific Cal3 nanobody with diagnostic and therapeutic potential(Hall *et al*., 2025). We harvested the Cal3 Nb from the periplasm of BL21 *E. coli* by osmotic shock, the established method for nanobody extraction(Pardon *et al*., 2014). However, the amount of Cal3 Nb in the periplasmic fractions and in the whole cell lysate was low (Fig. 1). Furthermore, most of Cal3 Nb was in the pellet fraction of an *E. coli* lysate that was separated by ultracentrifugation. These observations indicate that the majority of the Cal3 Nb is insoluble (Fig. 1).

**Fig. 1.**
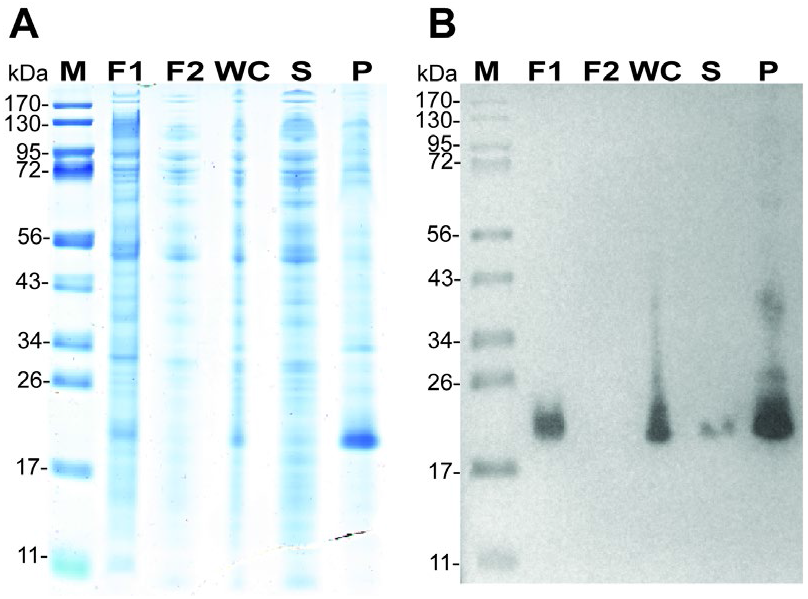
Expression and solubility of Cal3 Nb in *E. coli*. **(A)** SDS–PAGE analysis. Lanes: M, molecular weight marker; F1, periplasmic fraction 1; F2, periplasmic fraction 2; WC, whole-cell lysate; S, soluble fraction; P, pellet fraction. A total of ∼5 µg of protein was loaded in each lane. **(B)** Anti-FLAG Western blot analysis of the same samples following periplasmic extraction of Cal3 Nb and whole-cell lysis.

### Consensus Framework Mutations to Increase the Solubility of calreticulin Nanobodies

To improve the yield and the solubility of the Cal3 nanobody, we introduced 11 mutations to create the nanobody CALR-Nb02 (Fig. 2) to match the partial consensus framework as described by Dingus et al. (Dingus *et al*., 2022). The partial consensus sequence was defined as the amino acid sequence shared by at least 26 out of the 33 stable nanobodies analyzed in the study, and was strongly associated with improved solubility, intracellular stability and high yield of nanobodies. We also mutated the CDR2 cysteine 57 to serine to create the nanobody CALR-Nb03 to avoid the complications of potential intermolecular disulfide bonds between the free cysteines (Fig. 2). In addition, we removed the PelB signal sequence in the CALR nanobodies 02 and 03 to avoid their export to the periplasm, a likely yield-reducing step (Schlegel *et al*., 2013, Osgerby & Overton, 2023).

**Fig. 2.**
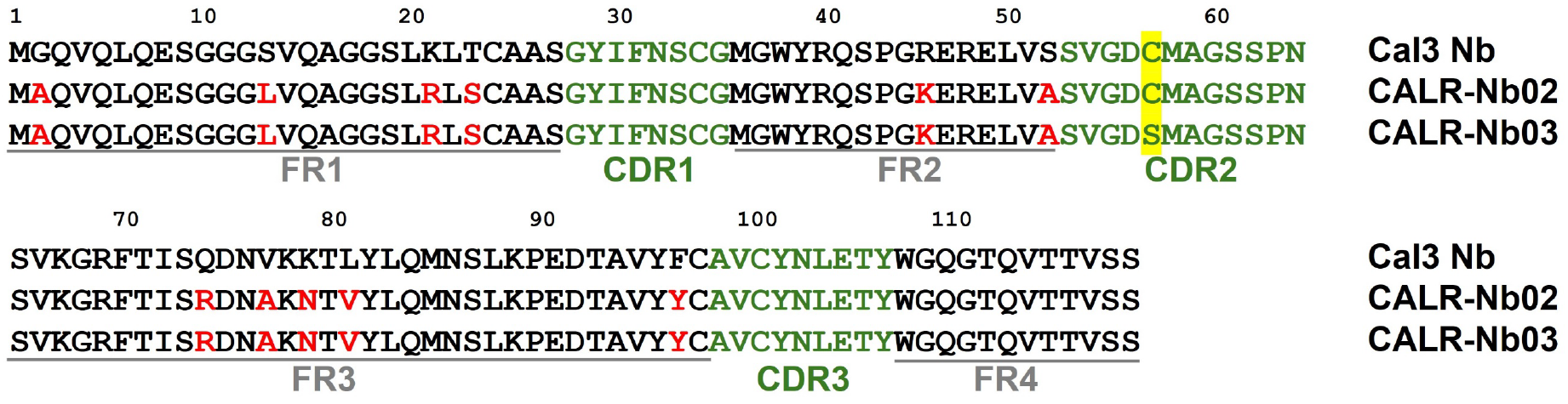
Sequence aligments of Cal3 Nb, CALR-Nb02 and CALR-Nb03. Alignment of the Cal3 nanobody sequence with the sequences of CALR-Nb02 and CALR-Nb03. Framework regions (FR) and complementarity-determining regions (CDR) are underlined in gray and shown in green letters, respectively. The framework mutations of the original Cal3 sequence to adapt the partial consensus sequence (Dingus *et al*., 2022) are shown in red. The cysteine 57 in CDR 2 for CALR-Nb02 was mutated to serine in CALR-Nb03 (highlighted in yellow).

### Comparison of *E. coli* Strains for Production of CALR Nanobodies

Since the formation of disulfide bonds is critical for the stability of the nanobody core (Mendoza *et al*., 2020) we chose *E. coli* Origami 2(DE3) as an expression host (Lopez-Cano *et al*., 2022). This strain lacks the glutathione reductase and thioredoxin reductase genes facilitating proper disulfide bond formation in the cytoplasm of *E. coli* (Bessette *et al*., 1999). To examine CALR-Nb02 nanobody production we expressed the *calr-nb02* gene under the control of the T7 promoter in *E. coli* Origami 2(DE3) and BL21 (DE3). Whole-cell lysates of both the *E. coli* Origami 2(DE3) and BL21 (DE3) strains show the highest expression levels of CALR-Nb02 four hours after induction (Fig. 3). Surprisingly, the CALR-Nb02 expression levels in the Origami 2 strain nearly approach that of the BL21 strain. Based on these expression results, we used *E. coli* Origami 2(DE3) for production of the CALR-Nb02 and CALR-Nb03 nanobody in the subsequent experiments because it enables disulfide bond formation in the cytoplasm.

**Fig. 3.**
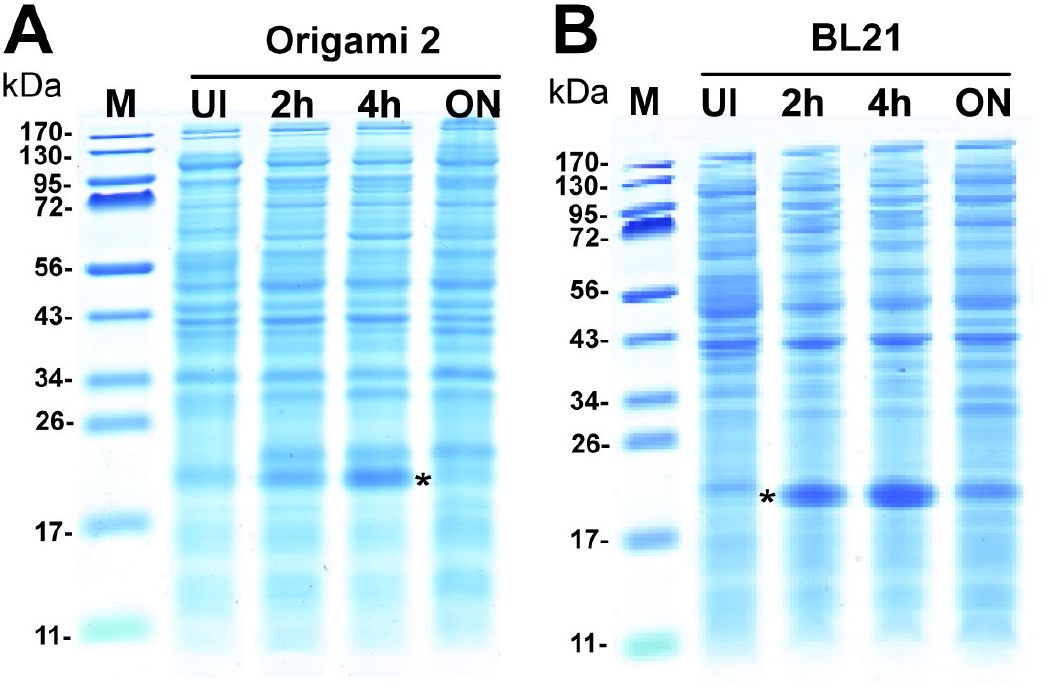
Whole cell lysate expression of CALR-Nb02 across *E. coli* strains. **(A)** SDS–PAGE analysis of whole cell lysates from Origami 2(DE3) *E. coli* expressing CALR-Nb02. For each strain, samples were collected before induction (UI) and at 2 hours, 4 hours, and overnight (ON) following induction. **(B)**SDS–PAGE analysis of whole cell lysates from BL21(DE3) *E. coli* expressing CALR-Nb02 collected under the same conditions. Molecular weight markers are shown in lane 1 of each gel. The position corresponding to CALR-Nb02 is marked with an asterisk.

### Framework Mutations and Cytoplasmic Expression Increase the Yield of CALR Nanobodies

The Cal3 Nb nanobody was produced in *E. coli* BL21 (DE3), while *E. coli* Origami 2(DE3) was used for CALR-Nb02 and CALR-Nb03 production. While Cal3 Nb was obtained by osmotic shock from the periplasm of *E. coli*, the CALR-Nb02 and CALR-Nb03 nanobodies were obtained by cell lysis from the cytoplasm of *E. coli*. The nanobodies were then purified by nickel affinity chromatography and subsequent size-exclusion chromatography (Fig. 4). SDS– PAGE analyses of the fractions showed increasing protein amounts from Cal3 Nb to CALR-Nb02 to CALR-Nb03 (Fig. 4). The yields of purified proteins for Cal3 Nb and CALR-Nb02 were 0.23 mg and 5.6 mg from one-liter cultures of *E. coli*. Thus, the framework mutations and cytoplasmic expression of CALR-Nb02 resulted in an approximately 240-fold increased protein yield. Importantly, most of the CALR-Nb02 protein was soluble and eluted as a monomer from the size exclusion column (not shown), providing further evidence for the value of the partial consensus framework for improving the yield of soluble nanobody as proposed previously (Dingus *et al*., 2022). The mutation of the single cysteine 57 increased the yield of purified nanobody CALR-Nb03 further by a factor of 3. This is likely due to the reduction of intermolecular disulfide bonds in CALR-Nb03 compared to CALR-Nb02. These results indicate that any free cysteine in addition to the conserved structural cysteine pairs in the nanobody core, are likely to reduce protein yield and solubility of nanobodies.

**Fig. 4.**
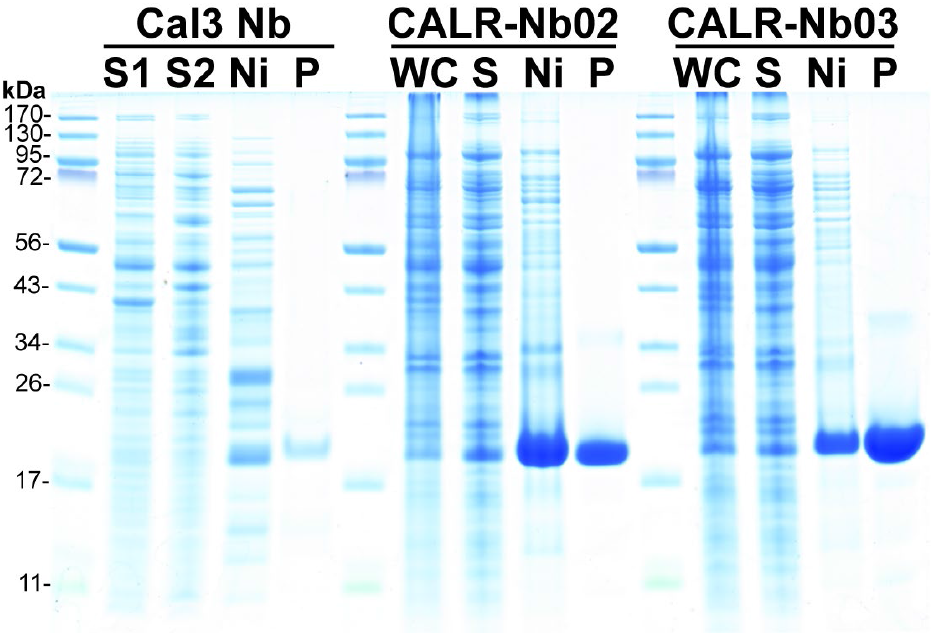
Purification of CALR nanobodies. Cal3 Nb, CALR-Nb02 and CALR-Nb03 were purified from *E. coli*. Samples were analyzed by Coomassie-stained 10% SDS-polyacrylamide gels. Lanes: S1, periplasmic extraction 1; S2, periplasmic extraction 2; WC, whole-cell lysate; S, soluble fraction; Ni, after Ni-affinity purification; P, purified protein after size exclusion chromatography. Nominally, 5 µg of each sample was loaded. However, unequal amounts of protein are visible in some lanes. Note, that due to low yield and concentration of Cal3 Nb, only 0.6 µg of protein was loaded.

### Framework Mutations Improve the Stability of CALR Nanobodies

To assess the thermal stability of the CALR nanobodies we used nano differential scanning fluorimetry, a dye-free technique that measures protein thermal stability by tracking changes in its intrinsic fluorescence. As the protein unfolds the fluorescence emission shifts from 330 nm to 350 nm due to the increased exposure of buried tryptophans to water (Kim *et al*., 2021). The traces of the fluorescence ratio (350 nm/330 nm) show a gradual increase without a clear transition for Cal3 Nb (Fig. 5A). By contrast, distinct unfolding transitions are visible for CALR-Nb02 and CALR-Nb03 (Fig. 5A). First-derivative analysis confirmed melting temperatures of 55.5 °C for CALR-Nb02 and 53.9 °C for CALR-Nb03, while no defined melting temperature could be assigned to Cal3 Nb (Fig. 5B). These results suggest that CALR-Nb02 and CALR-Nb03 are thermally stable, whereas Cal3 Nb lacks a clear transition, consistent with a heterogeneous sample likely resulting from aggregation and poor solubility.

**Fig. 5.**
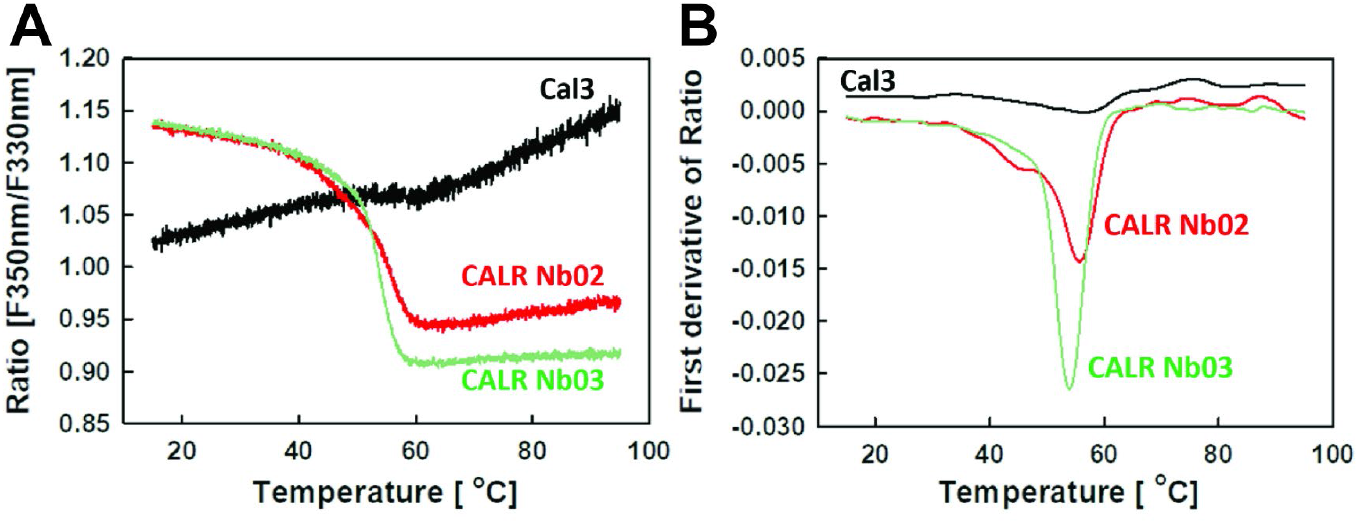
Nano differential scanning fluorimetry of CALR nanobodies. **(A)** Fluorescence intensity ratio (F350 nm/F330 nm). **(B)** First derivative of fluorescence intensity ratio. Capillaries were loaded with CALR Nb02 and CALR Nb03 at 0.5 mg/ml concentration. Concentration of Cal3 nanobodies was 0.1 mg/ml. Calculated inflection temperature of thermal unfolding, Tm, values were 36.6 °C and 76.2 °C for Cal3 Nb; 55.5 °C for CALR Nb02, and 53.9 °C for CALR Nb03. Values are mean from duplicate experiments.

### CALR-Nb02 Exhibits High Binding Affinity for Calreticulin

To examine whether the framework mutations in CALR-Nb02 affect the binding affinity for calreticulin, we used Bio-Layer Interferometry (BLI) (Calderin *et al*., 2025) and ELISA experiments. Both assays revealed strong binding of the CALR-Nb02 nanobody to calreticulin, with apparent affinities of 50.7 nM and 24.6 nM, respectively (Fig. 6). This represents a significant improvement over the previously reported Cal3 Nb affinity of approximately 140 nM (Hall *et al*., 2025). The apparent increased affinity of CALR-Nb02 is likely related to its improved solubility and folding, resulting in increased effective protein concentration compared to the Cal3 nanobody. The CALR-Nb03 nanobody did not show significant binding to calreticulin by ELISA (Fig. 6B), demonstrating that the cysteine 57 in the Cal3 and CALR-Nb02 nanobodies is essential for calreticulin binding.

**Fig. 6.**
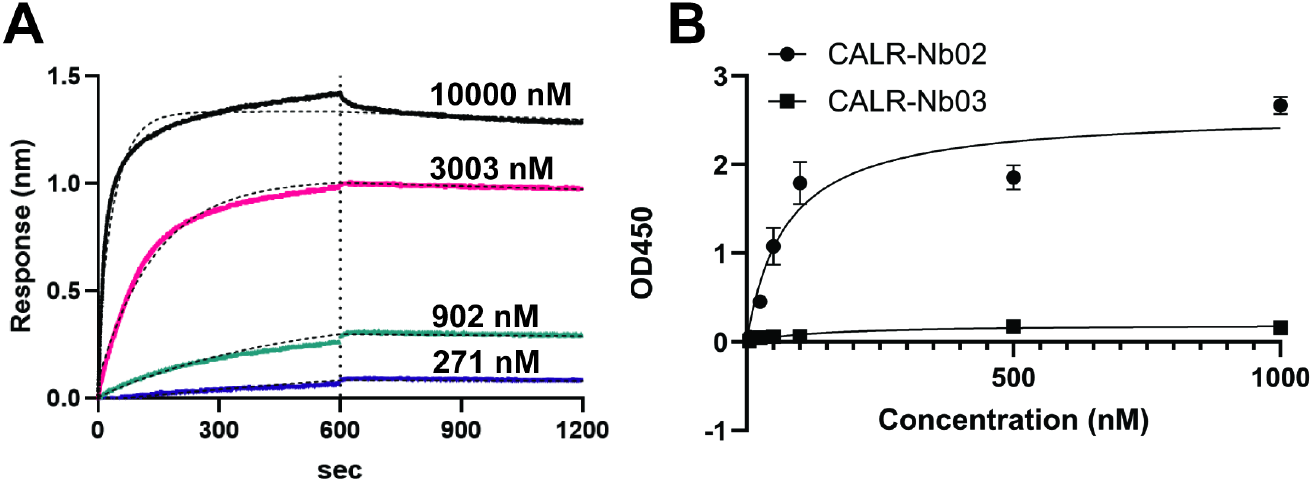
Affinity of CALR Nanobodies for Calreticulin (A) BLI sensorgrams of CALR-Nb02 binding to immobilized CALR at CALR-Nb02 concentrations ranging from 271 to 10,000 nM. A binding affiity of 50.7 nM was determined using a 1:1 binding model with global fit and steady-state analysis. **(B) ELISA of CALR-Nb02 (circles) and CALR-Nb03 (squares)** binding to calreticulin.

## DISCUSSION

In this study, we engineered a substantially improved calreticulin-specific nanobody CALR-Nb02 based on the previously reported Cal3 nanobody (Hall *et al*., 2025), which suffered from low solubility and protein yields. By combining partial consensus framework mutations (Dingus *et al*., 2022) with cytoplasmic expression in *E. coli*, the protein yield of the nanobody CALR-Nb02 was increased by 240-fold and its solubility and thermal stability were substantially improved compared with Cal3 Nb. This improvements also resulted in stronger calreticulin binding with a K_D_ of 25-50 nM which is similar similar to that of nanobodies derived from naïve libraries whose dissociation constants range from 10 nM to 150 nM (Oloketuyi *et al*., 2021). These results show that CALR-Nb02 is a much better calreticulin-binding reagent than the parent Cal3 nanobody.

Our results also suggest that protein yields obtained by Cal3 Nb are limited by poor solubility due to misfolded protein which is degraded in *E. coli* and/or lost during protein purification due to aggregation (Betton *et al*., 1998). This may be a consequence of the original engineering strategy, in which the calreticulin-binding CDR2 (Fig. 2) was separately evolved and grafted onto the otherwise unchanged HER2-specific nanobody 2Rs15d (Hall *et al*., 2025). Although CDR grafting can transfer binding specificity to other nanobodies, CDR loops often do not function independently from the scaffold that supports them (Gil & Schrum, 2013). This is because the relatively long CDR2 loop sequences can strongly influence nanobody folding and stability, due to their close proximity to the nanobody core (Perchiacca & Tessier, 2012). Consistent with this notion, grafting only one CDR from one antibody domain onto another often reduces stability relative to the parental domains, whereas grafting all three CDRs results more frequently in stable scaffolds (Perchiacca & Tessier, 2012). The strong improvements observed for CALR-Nb02 compared to the Cal3 nanobody demonstrate that framework mutations not only can rescue poorly soluble, grafted nanobodies but also reveal a novel strategy for grafting stable and functional nanobodies by transferring a single complimentary determining region with an altered binding affinity to already optimized nanobodies. It should be noted that the likely lack of interactions of amino acids of the grafted CDR with those of other CDRs and the framework in a grafted nanobody likely decrease its stability as shown for the below average thermal stability of CALR-Nb02. Taken together, our results show that the framework mutations as defined by Dingus et al. (Dingus *et al*., 2022) drastically improve the stability of the nanobody core enabling nanobodies to tolerate the lack of beneficial interactions between a grafted CDR2 and the nanobody core. Such a strategy may also work for other small protein binders such as affibodies (Blanchard *et al*., 2023, Crook *et al*., 2020).

While the stability of CALR-Nb02 is drastically improved compared to the Cal3 nanobody, its melting temperature of ∼55 °C is lower than the average experimental melting temperatures of natural or evolved nanobodies ranging between 60 °C and 80 °C with an overall median of ∼67 °C (Valdes-Tresanco *et al*., 2023, Alvarez & Dean, 2024) (Muyldermans, 2013). The lower melting temperature of CALR-Nb02 likely reflects the lack of stabilizing interactions with the CDR2 and the core of CALR-Nb02 (Perchiacca & Tessier, 2012) exposing another limitation of the grafting strategy. However, if thermal stability is an important feature of a particular nanobody application, alternative stability engineering approaches have been described (Kunz *et al*., 2017, Kunz *et al*., 2019) in addition to the framework mutations utilized in our study (Dingus *et al*., 2022).

Interestingly, the CDR2 of CALR Nb02 contains an essential single cysteine at position 57. Surface-accessible single cysteines such as C57 of CALR Nb02 are prone to form intermolecular disulfide bonds. This can result in oligomer formation during protein folding reducing the yield or dimerization of folded proteins under oxidizing conditions (Kang *et al*., 2025). Indeed, we observed both increased dimer formation for Ni(II) affinity purified CALR Nb02 vs Nb03 (Fig. 4) and reduced yields for CALR Nb02 vs Nb03. These observations are consistent with other proteins with unpaired or noncanonical cysteines which were found to reduce the yield of soluble protein due to heterogeneous disulfide bonding (Schumacher *et al*., 2018). The similar stability and solubility of both CALR nanobodies indicate that C57 directly participates in calreticulin recognition or stabilizes the local CDR conformation required for binding.

Monoclonal antibodies are widely used in modern medicine (Burton *et al*., 2012, Caskey *et al*., 2019, Corti et al., 2021, Castelli et al., 2019). However, major advantages of nanobodies over monoclonal antibodies are their small size, high stability and cheap production (Muyldermans, 2021, Abdellatif *et al*., 2026). In this study we have improved the solubility, stability, yield and target affinity of the original grafted Cal3 nanobody by engineering the nanobody core. These properties make CALR-Nb02 suitable for applications that require high yield protein production and specific recognition of calreticulin as a biomarker for multiple cancers that could be utilized for tumor detection, prognosis, and treatment response (Fucikova *et al*., 2021, Kepp *et al*., 2020). Our improved calreticulin-specific nanobody is, therefore, a useful tool for both diagnostic assay development, including blood-based calreticulin testing, and future theragnostic strategies targeting calreticulin-positive tumors.

## EXPERIMENTAL PROCEDURES

### Strains and Plasmids

All bacterial strains and plasmids used in this study are listed in Table S1. *Escherichia coli* DH5α was used for all cloning procedures. *E. coli* BL21(DE3) and Origami 2(DE3) were used for the expression of recombinant nanobodies. The *E. coli* strains were routinely grown in Lysogenic Broth (LB) medium at 37 °C unless indicated otherwise. The following antibiotics were used when required at the following concentrations: ampicillin (100 µg/mL for *E. coli*), tetracycline (12.5 µg/mL for Origami 2(DE3) *E. coli*), streptomycin (50 µg/mL for Origami 2(DE3) *E. coli*).

### Expression and Purification of nanobodies from *E. coli*

Cal3 Nb was expressed in the periplasm of *E. coli*. The plasmid for Cal3 Nb was transformed into BL21(DE3) *E. coli* cells. Cells were cultured in 1 L supplemented with 0.1% glucose and carbenicillin and incubated at 37 °C at 200 rpm. When the OD600 reached 0.5, the cells were induced with 0.1mM Isopropyl β-D-1 thiogalactopyranoside (IPTG) and incubated at 37 °C and 200 rpm for 4 hours. Cultures were harvested by centrifugation at 4000 × g for 20 minutes, followed by resuspension of the cell pellets in phosphate buffered saline (PBS) and subsequent centrifugation at 3200 × g for 10 minutes. Cell pellets were stored in -20 °C until further use. Cell pellet was resuspended in SET buffer (50 mM Tris HCL, 1 mM EDTA, 20% sucrose, pH 7.5) in a 1 to 5 (w:v) ratio. The suspension was incubated for 10 minutes at 4 °C and 200 rpm then centrifuged for 15 minutes at 8,000 x g and 4 °C. The supernatant was removed and stored at 4 °C, and the remaining cell pellet was resuspended in chilled puirfied water (Millipore) in a 1 to 5 (w:v) ratio and then incubated on ice for 5 minutes. The cell suspension was centrifuged for 15 minutes at 8,000 x g at 4 °C and the supernatant was collected and stored at 4 °C. Nanobody was further purified by Ni-Affinity chromatography (Marvelgent Biosciences 11-0227-050) and size exclusion chromatography (Bio-Rad Sdex 75).

CALR-Nb02 plasmid was transformed into BL21(DE3) and Origami 2(DE3) *E. coli* cells. All cells were cultured in 1 L LB medium supplemented with 0.1% glucose and carbenicillin at 37 °C at 200 rpm. Origami 2(DE3) cultures were additionally supplemented with tetracycline and streptomycin. When the OD600 reached 0.5, the cells were induced with 0.1 mM IPTG and incubated at 37 °C and 200 rpm. Aliquots of each culture were taken before induction, 2 hours after induction, 4 hours after induction, and after overnight induction. Cells were harvested by centrifugation at 4000 × g for 20 minutes, followed by resuspension of the cell pellets in PBS and subsequent centrifugation at 3200 × g for 10 minutes. Cells were resuspended in lysis buffer (PBS, protease inhibitor cocktail, benzonase) in a 1 to 15 (w:v) ratio and lysed by sonication. Protein concentration of whole cell lysates was determined using the Bradford Reagent. 15 µg of each sample were mixed with loading dye and 1% beta-mercaptoethanol, incubated at 100 °C for 3 minutes, and loaded onto acrylamide gel for SDS-PAGE analysis.

CALR-Nb02 and CALR-Nb03 were expressed in the cytoplasm of *E. coli* Origami 2(DE3). The bacteria were cultured in 1 L supplemented with 0.1% glucose, tetracycline, streptomycin and carbenicillin at 37 °C at 200 rpm. When the OD600 reached 0.5, the cells were induced with 0.1 mM IPTG and incubated at 37 °C and 200 rpm for 4 hours. Cultures were harvested by centrifugation at 4000 × g for 20 minutes, followed by resuspension of the cell pellets in PBS and subsequent centrifugation at 3200 × g for 10 minutes. Cell pellets were resuspended in lysis buffer (PBS, protease inhibitor cocktail, benzonase) in a 1 to 15 (w:v) ratio and the cells were lysed by sonication. The soluble fraction was isolated by high-speed centrifugation for 20 minutes at 20,000 x g and 4 °C. Nanobodies were further purified by Ni-Affinity chromatography (Marvelgent Biosciences 11-0227-050) and size exclusion chromatography (Bio-Rad Sdex 75).

### Solubility Analysis of Nanobodies

Cell pellets were resuspended in lysis buffer (PBS, protease inhibitor cocktail, benzonase) in a 1 to 15 (w:v) ratio. The cells were lysed by sonication. The soluble fraction was isolated by high speed centrifugation for 20 minutes at 20,000 x g and 4 °C. 5 µg of each sample were mixed with loading dye and 1% beta-mercaptoethanol, incubated at 100 °C for 3 minutes, loaded on acrylamide gel for SDS-PAGE analysis and transferred to PVDF for western blot. Anti-FLAG HRP antibody (Sigma Aldrich A8592) was used for western blot.

### Nanobody Sequence Design and Cloning

The amino acid sequence of Cal3 Nb was obtained from the preprint (Hall *et al*., 2025). Framework residues were modified at selected positions based on reported stabilization mutations to construct CALR-Nb02. The CALR-Nb02 sequence was codon optimized for *E. coli* expression, synthesized d*e novo* (GenScript), and cloned into the pET22b expression vector using HindIII and NdeI restriction enzymes. CALR-Nb02 omitted the PelB leader Sequence from Cal3 Nb but features a C-terminal 6x histidine tag and FLAG tag. CALR-Nb03 was constructed by mutating the CDR2 cysteine at position 143 in CALR-Nb02 to serine using T7 forward and reverse primers and a mutagenesis primer (CAAGTGTTGGGGATTCTATGGCGGG).

### Nano Differential Scanning Fluorimetry

NanoDSF was performed using Prometheus NT.48 instrument (NanoTemper Technologies, Munchen, Germany). Purified protein samples in 100 mM HEPES, 2 mM TCEP, pH 7.2 buffer were loaded in standard nanoDSF capillaries from NanoTemper Technologies. Protein concentrations were 0.5 mg/ml for CALR-Nb02 and CALR-Nb03, and 0.1 mg/ml for Cal3 Nb. Capillaries were exposed to temperatures from 15 °C to 95 °C with thermal ramping rate of 2 °C/min for the duration of 40 minutes. Fluorescence emission from tryptophan after UV excitation at 280 nm was collected at 330 nm and 350 nm with dual-UV detector. Inflection temperature of thermal unfolding, Tm, was calculated by PR.ThermControl software (NanoTemper Technologies, Munchen, Germany). Data then were exported into Excel (Microsoft) and graphs were prepared using SigmaPlot 11.0 (Systat Software, Inc).

### Binding affinity of nanobodies by Biolayer Interferometry (BLI)

Biolayer interferometry (BLI) was performed using an Octet RED (ForteBio) controlled by Octet DataAnalysis software (version 9.0.0.14). Streptavidin (SA) biosensors (Sartorius, 18-5019) were hydrated in TBST for at least 10 minutes before use. For ligand immobilization, biotinylated CALR was diluted to 10 µg/mL in TBST and loaded onto the biosensors for 600 s at 1000 rpm. After loading, biosensors were washed in TBST to remove unbound ligand. Binding kinetics were assessed by exposing the functionalized biosensors to a series of CALR-Nb02 concentrations ranging from 23.8 nM to 22.7 µM in a 3.33-fold dilution series. Association and dissociation were monitored for 300 s each at room temperature (RT). A reference biosensor was used to correct for nonspecific binding. Data were analyzed using Octet DataAnalysis software (version 9.0.0.14), and kinetic parameters (k_a_, kd, and KD) were determined using a 1:1 binding model with global fit and steady-state analysis. Data were baseline corrected using reference well subtraction, and negative controls included a buffer-only reference well.

### Binding affinity of nanobodies by the Enzyme-Linked Immunosorbent Assay (ELISA)

Recombinant Calreticulin (Sino Biological 13539-H08H) was diluted in 0.1 M sodium bicarbonate buffer at pH 9.5 and 100 ng were immobilized per well on Maxisorp 96-well plates (ThermoFisher). Wells were blocked overnight with 3% milk in PBS at 4 °C. Nanobody at concentrations ranging from 1 nM to 1000 nM was added to the wells in triplicate and incubated for 1 hour at room temperature with shaking. Wells were washed five times with 300 µL of 1X TBST. An anti-FLAG horseradish peroxidase conjugated antibody (Sigma Aldrich A8592) was diluted 1 to 1000 and added to each well followed by 1 hour incubation at room temperature with shaking. Wells were washed five times with 1X TBST. TMB (Millipore Sigma) was added to each well and incubated at room temperature for 15 minutes with shaking and covered in foil. The reaction was quenched by addition of 0.2 M sulfuric acid. Absorbance at 450 nm was measured using a BioTek plate reader. Binding curves were generated using GraphPad Prism version 10.4.1 by nonlinear regression analysis using a one site binding model.

## Acknowledgments

We thank Southern Research for access to Octet RED for the biolayer interferometry experiments and Dr. Champy Deivanayagam for access to Prometheus NT48 for nano differential scanning fluorimetry. This work was supported by a grant from the UAB O’Neal Comprehensive Cancer Center to MN and BL and by a Triton Endowment Fund (UAB) to MN. LM was supported by the Innovate Alabama Network Designation Grant and UAB’s Clinical Diagnostics Sciences Department.

## Abbreviations

CALR: calreticulin
OPOE: *n*-octylpolyethylene oxide
LDAO: lauryldimethylamine oxide
C_12_E_8_: octaethylene glycol monododecyl ether

## Data Availability Statement

The data underlying this article will be shared on reasonable request to the corresponding author.

